# Neuronal populations in the occipital cortex of the blind synchronize to the temporal dynamics of speech

**DOI:** 10.1101/186338

**Authors:** Markus J. van Ackeren, Francesca Barbero, Stefania Mattioni, Roberto Bottini, Olivier Collignon

## Abstract

The occipital cortex of early blind individuals (EB) activates during speech processing, challenging the notion of a hard-wired neurobiology of language. But, at what stage of speech processing do occipital regions participate in EB?Here we demonstrate that parieto-occipital regions in EB enhance their synchronization to acoustic fluctuations in human speech in the theta-range (corresponding to syllabic rate), irrespective of speech intelligibility. Crucially, enhanced synchronization to the intelligibility of speech was selectively observed in primary visual cortex in EB, suggesting that this region is at the interface between speech perception and comprehension. Moreover, EB showed overall enhanced functional connectivity between temporal and occipital cortices sensitive to speech intelligibility and altered directionality when compared to the sighted group. These findings suggest that the occipital cortex of the blind adopts an architecture allowing the tracking of speech material, and therefore does not fully abstract from the reorganized sensory inputs it receives.

## Introduction

The human cortex comprises a number of specialized units, functionally tuned to specific types of information. How this functional architecture emerges, persists, and develops throughout a person’s life are among the most challenging and exciting questions in neuroscience research. Although there is little debate that both genetic and environmental influences affect brain development, it is currently unknown how these two factors jointly shape the functional architecture ofthe cortex. A key topic in this debate is the organization of the human language system. Language is commonly thought to engage a well-known network of regions around the lateral sulcus. The consistency of this functional mapping across individuals, as well as its presence early in development are remarkable, and often used as an argument that the neurobiological organization of the human language system is the result of innate constraints (Dehaene-Lambertz et al., 2006; Berwick et al., 2013). Does the existence of a highly consistent set of regions for language acquisition and processing imply that this network is “hardwired” and immutable to change by experience?Strong nativist proposals for linguistic innateness leave little room for plasticity due to experience (Bates, 1999), with for instance the proposition that we should conceive “the growth of language as analogous to the development of a bodily organ” (Chomsky, 1975, p. 11). However, studies in infants born with extensive damage tothe typical brain’s language areas may develop normal language abilities, demonstrating that the language network is subject to reorganization (Bates, 2005). Perhaps the most intriguing demonstration that the neurobiology of language is susceptible to change due to experience comes from studies in congenitally blind individuals showing functional selectivity to language in primary and secondary ‘visual’ areas (Roder et al., 2002; Burton, 2003; Amedi et al., 2004; Bedny et al., 2011; Arnaud et al., 2013). Such reorganization of the language network is particularly fascinating since it arises in a system with no injuries ofthe core language network (Bates, 2005; Atilgan et al., 2017).

However, the level of speech representation at which the occipital cortex of the early blind participates remains poorly understood. Speech comprehension requires that the brain extract meaning from the acoustic features of sounds(de Heer et al., 2017). Although several neuroimaging studies have yielded valuable insights about the processing of speech in early blind adults (Arnaud et al., 2013; Bedny et al., 2011; Burton, 2003; Lane et al., 2015; Röder et al., 2002)and infants (Bedny et al., 2015), these methods do not adequately capture the fast and continuous nature of speech processing. Because speech unfolds over time, understanding spoken language rely on the ability to track the incoming acoustic signal in near real-time (Peelle and Davis, 2012). Indeed, speech is a fluctuating acoustic signal that rhythmically excites neuronal populations in the brain (Poeppel et al., 2008; Gross et al., 2013; Peelle et al., 2013). Several studies have demonstrated that neuronal populations in auditory areas entrain to the acoustic fluctuations that are present inhuman speech around the syllabic rate (Luo and Poeppel, 2007; Kayser et al., 2009; Szymanski et al., 2011; Zoefel and VanRullen, 2015). It has therefore been suggested that entrainment reflects a key mechanism underlying hearing, for example, by facilitating the parsing of individual syllables through adjusting the sensory gain relative to fluctuations in the acoustic energy (Giraud and Poeppel, 2012; Peelle and Davis, 2012; Ding and Simon, 2014). Crucially, because some regions that track the specific acoustic rhythm of speech are sensitive to speech intelligibility, neural phase-locking is not solely driven by changes in the acoustic cue of the auditory stimuli, but also reflects cortical encoding and processing of the auditory signal (Peelle et al., 2013; Ding and Simon, 2014). Speech tracking is therefore an invaluable tool to probe regions interfacing speech perception and comprehension Poeppel et al., 2008; Gross et al., 2013; Peelle et al., 2013).

Does the occipital cortex of early blind people synchronizes to speech rhythm? Is this putative phase-locking of neural activity to speech influenced by its understandability? Addressing those questions would provide novel insights onto the functional organization of speech processing rooted in the occipital cortex of early blind people. In the current study, we investigated if neuronal populations in blind occipital cortex synchronize to rhythmic dynamics of speech. This tracking is demonstrated by relating the amplitude fluctuations in speech to electromagnetic dynamics recorded from the participant’s brain. To this end, we recorded the brain activity of a group of early blind (EB; n=17) and sighted individuals (SI; n=16) with Magnetoencephalography (MEG), while participants listened to short narrations from audiobooks. If the occipital cortex of the blind entrains to speech rhythms, this will support the idea that this region processes low-level acoustic features relevant for language understanding. We further tested whether the putative phase locking of occipital responses in early blind people benefits from linguistic information or only relates to acoustic information. To separate linguistic and acoustic processes we relied on a noise-vocoding manipulation that spectrally distorted the speech signal in order to gradually impair intelligibility but systematically preserve the slow amplitude fluctuations responsible for speech rhythm (Shannon et al. 1995; Peelle et al., 2012).

Furthermore, going beyond differences in the local encoding of speech rhythm in the occipital cortex, we also investigated whether early blindness alters the connectivity between occipital and temporal regions sensitive to speech comprehension.

## Results

### Story comprehension

Participants listened to either natural speech segments (nat-condition) or to altered (using vocoding) versions of these segments (see material and methods for details). In the 8-channel vocoded condition, the voice of the speaker is highly distorted but intelligibility is unperturbed. In contrast, in the 1-channel vocoded condition, speech is entirely unintelligible. After listening to each speech segment, participants were provided with a short statement about the segment, and asked to indicate whether the statement was true or false. Behavioural performance on these comprehension statements was analysed using linear mixed-effects models with maximum likelihood estimation. The method is a linear regression that takes into account dependencies in the data, as present in repeated measures designs. Blindness and Intelligibility were included as fixed effects, while subject was modelled as a random effect. Intelligibility was nested in subjects. Intelligibility had a significant effect on story comprehension (*X*(2)=110.7, *p*<001). The effect of blindness and the interaction between intelligibility and blindness were non-significant (*X*(1)=1.14, *p*=.286 and *X*(2)=.24, *p*=.889). Orthogonal contrasts demonstrated that speech comprehension was stronger in the nat and 8-channel condition versus the 1-channel condition (b=.78, *t*(62)=14.43, *p* <001, r=.88). There was no difference between the nat and the 8-channel condition (b=.02, *t*(62), 1.39, *p*=.17). Thus, speech intelligibility was reduced in the 1-channel, but not the 8-channel vocoded condition (Fig 1D). The lack of effect for the factor blindness suggests that there is no evidence for a potential difference in comprehension, attention or motivation between groups. Descriptive statistics of the group, and condition means are depicted in Table 1.

**Table 1.** Proportion of correct responses for the story comprehension questions. Depicted are the number of subjects (N) in each group, the mean (M), and the standard error of the mean (SE) for each of the three conditions (Natural, 8-channel, 1-channel).

| Group | N | <u>Natural</u> |  | <u>8-channel</u> |  | <u>1-channel</u> |  |
| --- | --- | --- | --- | --- | --- | --- | --- |
|  |  | M | SE | M | SE | M | SE |
| EB | 17 | 0.899 | 0.019 | 0.853 | 0.031 | 0.559 | 0.027 |
| SI | 15 | 0.914 | 0.020 | 0.890 | 0.025 | 0.576 | 0.034 |

### Tracking intelligible speech in blind and sighted temporal cortex

The overall coherence spectrum, which highlights the relationship between the amplitude envelope of speech and the signal recorded from the brain, was maximal over temporal sensors between 6-7 Hz (Fig 1E). This first analysis was performed on the combined dataset, and hence does not cause the risk of a bias or circularity for subsequent analyses, targeting group differences. The peak in the current study is slightly higher than what has been reported in previous studies (Gross et al., 2013). A likely explanation for this shift is the difference in syllabic rate between English (~6.2Hz), used in previous studies, and Italian (~7Hz) (Pellegrino et al., 2011). The syllabic rate is the main carrier of amplitude fluctuations in speech, and thus most prone to reset oscillatory activity.

**Fig. 1.**
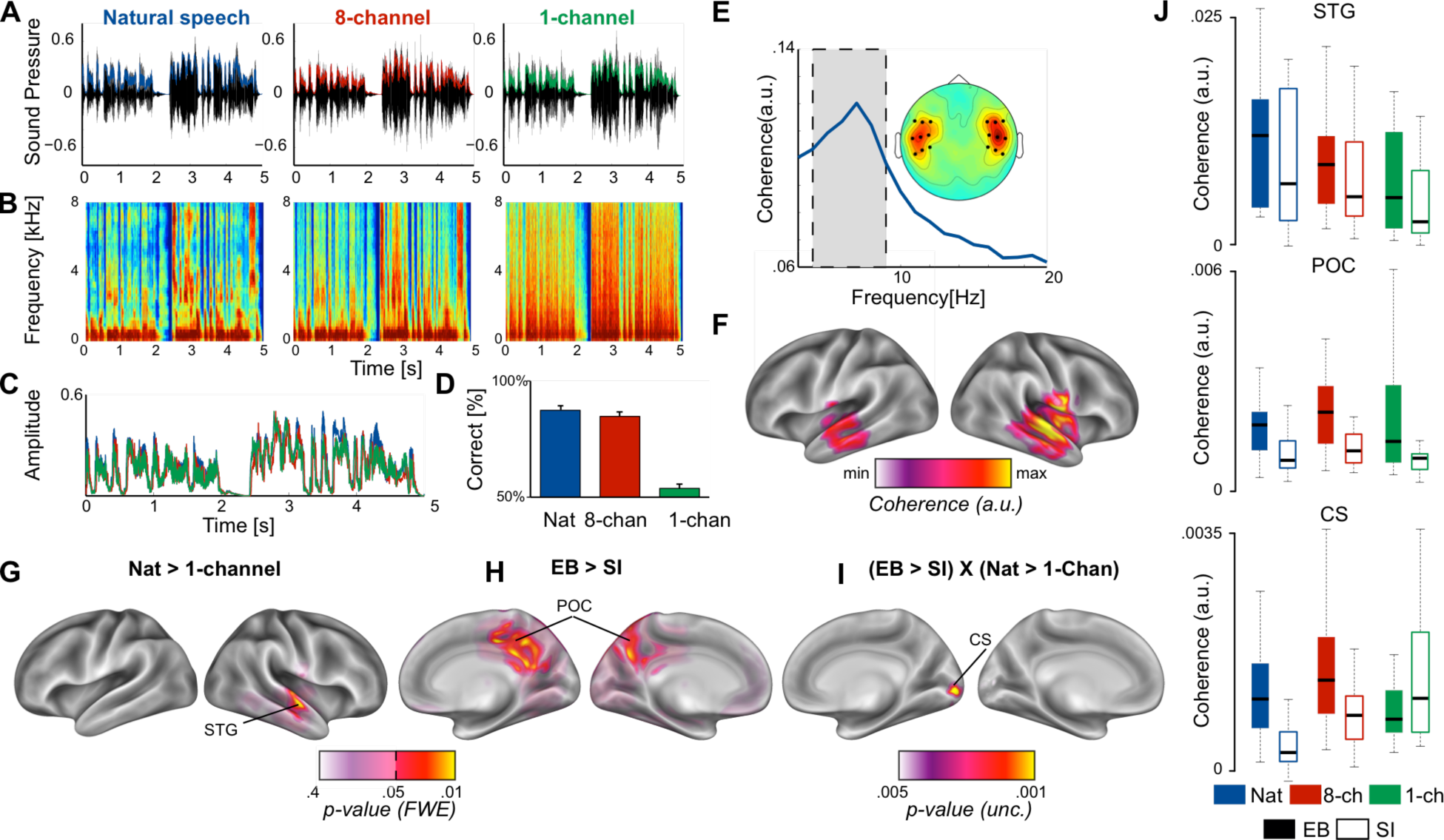
Cerebro-acoustic coherence in EB and SI. (A) Segments from each condition (nat, 8-channel, and 1-channel) in the time domain. Overlaid is the amplitude envelope of each condition. (B) Spectrogram of the speech samples from (A) The effect of spectral vocoding on the fine spectral detail in all three conditions. (C) The amplitude envelopes of the samples for each condition superimposed are highly similar, despite distortions in the fine spectral detail. (D) Behavioural performance on the comprehension statements reveals that comprehension is unperturbed in the nat and 8-channel conditions, whereas the 1-channel conditions elicits chance performance. (E) Coherence spectrogram extracted from bilateral temporal sensors reveals a peak in cerebro-acoustic coherence at 7Hz, across groups and conditions. The shaded area depicts the frequency range of interest in the current study. The topography shows the spatial extent of coherence values in the range 4-9Hz. Enhanced coherence is observed selectively in bilateral temporal sensors. (F) Source reconstruction of the raw coherence confirms that cerebro-acoustic coherence is strongest in the vicinity of auditory cortex, bilaterally, extending into superior and middle temporal gyrus. (G) Statistically thresholded maps for the contrast between natural and 1-channel vocoded speech show an effect of intelligibility (*p*<.05, FWE-corrected) in right STG. (H) Enhanced envelope tracking is observed for EB versus SI in a bilateral parieto-occipital network network along the medial wall centred on Precuneus (*p*<.05, FWE-corrected). (I) The statistical map shows the interaction effect between blindness and intelligibility: Early blind individuals show enhanced synchronization during intelligible (nat) versus non-intelligible speech (1-channel)as compared to SI in right calcarine sulcus (CS) (*p*<.005, uncorrected). (J) Boxplots for three regions identified in the whole brain analysis (top panel: STG; middle panel: parieto-occipital cortex; bottom panel: calcarine sulcus).

To optimally capture the temporal scale of cerebro-acoustic coherence effects (6-7Hz), and achieve a robust source estimate, source reconstruction was performed on two separate frequency windows (6±2, and 7±2Hz) for each subject and condition. The two source images were averaged for subsequent analysis, yielding a single image representing the frequency range 4-9Hz. The source-reconstructed coherence at the frequency of interest confirmed that cerebro-acoustic coherence was strongest in bilateral temporal lobes including primary auditory cortex. (Fig 1F).

To test whether envelope tracking in the current study is modulated by intelligibility, we compared the coherence maps for the intelligible (nat), versus non-intelligible (1-channel) condition, in all groups combined (i.e., EB and SI), with dependent-samples permutation t-tests in SPM. The resulting statistical map (Fig 1G, *p*<.05, FWE-corrected), revealed a cluster in right superior, and middle temporal cortex (STG, MTG) where synchronization to the envelope of speech was stronger when participants had access to the content of the story.

### Speech tracking in blind parieto-occipital cortex

Whether early blind individuals recruit additional neural substrate for tracking the envelope of speech was tested using independent-samples permutation t-tests in SPM contrasting coherence maps between EB and SI for all three conditions combined. The statistical maps (Fig. 1H, *p*<.05, FWE-corrected) revealed enhanced coherence in EB versus SI in parieto-occipital cortex along the medial wall, centered on bilateral Precuneus, branching more extensively into the right hemisphere (see Table 2). These results demonstrate that neuronal populations in blind parieto-occipital cortex synchronize to the temporal dynamics in speech.

**Table 2.** MNI coordinates and p-values for the three contrasts tested with TFCE

|  |  | <u>MNI-coordinates (mm)</u> |  |  |
| --- | --- | --- | --- | --- |
|  | *p-value* (FWE-cor) | x | y | z |
| Intelligibility |  |  |  |  |
| Superior Temporal Gyrus | 0.027 | 64 | -16 | 0 |
| Blindness |  |  |  |  |
| Posterior Cingulate | 0.030 | 8 | -40 | 40 |
| Postcentral sulcus | 0.033 | 16 | -32 | 64 |
| Precuneus | 0.033 | -8 | -64 | 48 |
| Blindness X Intelligibility |  |  |  |  |
| Calcarine sulcus | 0.007 | 16 | -88 | 0 |

Finally, to investigate if, and where envelope tracking is sensitive to top-down predictions during intelligible speech comprehension, we subtracted the unintelligible (1-channel) from the intelligible (nat) condition, and computed independent-samples t-tests between groups. As highlighted in the introduction, sensitivity to language is most surprising in early ‘visual’ areas of blind occipital cortex. Especially the role of the area around Calcarine sulcus (V1) is currently debated (Burton et al., 2002; Roder et al., 2002; Amedi et al., 2003). As there is no prior on the potential effect size for this contrast we restricted the analysis to the area around primary visual cortex. The search volume was constructed from four 10mm spheres around coordinates in bilateral Calcarine sulcus ([-7, −81, −3]; [-6, −85, 4]; [12, −87, 0]; [10, −84, 6]). The coordinates were extracted from a previous study on speech comprehension in EB (Burton et al., 2002). The resulting mask also extends into regions highlighted in similar studies by other groups (Roder et al., 2002; Bedny et al., 2011).

A significant effect for the interaction between blindness and intelligibility [(nat_EB_ – 1-channel_EB_) - (nat_SI_ – 1-channel_SI_)] was observed in right Calcarine sulcus. To explore the spatial specificity of the effect, we show the whole-brain statistical map for the interaction between intelligibility and blindness at a more liberal threshold (*p*<.005, uncorrected) in Fig 1I. The figure shows that intelligible speech selectively engages the area around the right calcarine sulcus in the blind versus sighted. Specifically, the region corresponding to right ‘visual’ showed enhanced sensitivity to intelligible speech in EB versus SI (*p*<.05, FWE-corrected). The analysis contrasting thetwo intelligible conditions (nat, 8-channel), did however not yield a significant effect in the EB, suggesting that the effect is not likely driven by the low-level degrading of the stimulus alone, but rather the intelligibility of the speech segment. Follow-up post-hoc comparisons between the two groups revealed that coherence was stronger during the intelligible condition for EB versus SI (t(30)=3.09, *p*=.004), but not during the unintelligible condition (t(30)=-1.08, *p*=.29). Nevertheless, inspection of the data suggests that the overall interaction effect was influenced by high coherence during the unintelligible condition in the SI group. For illustration purposes, cerebro-acoustic coherence from functional peak locations for intelligibility (across groups) in STG, blindness in POC (Fig 1G), and the interaction, are represented as boxplots in Fig 1J.

### Changes in occipito-temporal connectivity in the early blind

To further investigate whether cerebro-acoustic peaks in CS, and STG, which are also sensitive to intelligible speech in EB, are indicative of a more general re-organization of the network, we applied functional connectivity analysis. Statistical analysis of the connectivity estimates was performed on the mean phase-locking value in the theta (4-8Hz) range

Using linear mixed-effects models, blindness (EB, SI), intelligibility (nat,1-channel), and the interaction between blindness and intelligibility were added to the model in a hierarchical fashion. Blindness and intelligibility were modelled as fixed effects, while subject was a random effect. Intelligibility was nested in subject. A main effect was observed only for blindness (*X*(1)=4.32, *p*=.038). That is, connectivity was overall stronger for EB versus SI (Fig 2 A-B). The main effect of intelligibility, and the interaction between intelligibility and blindness were non significant (*p*=.241, and *p*=.716 respectively)

**Fig. 2.**
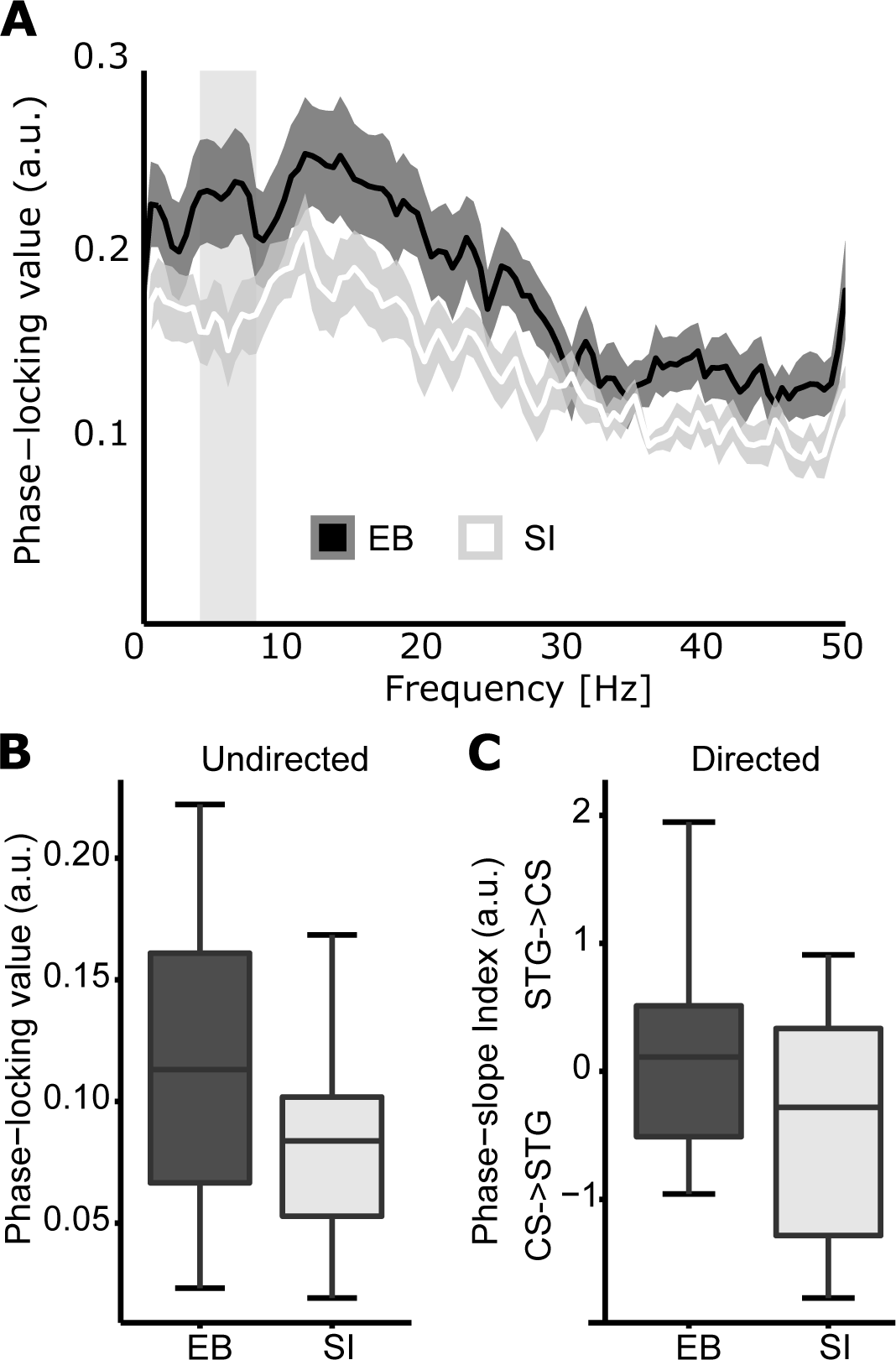
Occipital-temporal connectivity in EB and SI. (A) Spectra illustrate the phase locking between CS and STG in EB (dark curve), and SI (light curve). Shaded areas illustrate the SEM. A difference between the curves is observed in the theta range. (B) Boxplots depict the mean phase locking between CS and STG for EB (black), and SI (white) in the theta range. Connectivity is enhanced in EB versus SI. (C)Directional connectivity analysis using the phase-slope index (PSI). Positive values suggest enhanced directionality from STG to CS, and negative values represent enhanced connectivity in the opposite direction. He boxplots highlight that SI show a strong feed-forward drive from CS to STG, whereas blind individuals show a more balanced pattern, with a non-significant trend in the opposite direction.

Subsequently, linear mixed-effects models were applied to test for the effects blindness, and intelligibility on the directional information flow (PSI) between CS and STG. The fixed and random effects structure was the same as described in the previous analysis. Here, only the main effect of blindness was significant (*X*(1)=4.54, *p*=.033). That is, the directional information flow differs between groups. The effect of intelligibility and the interaction between intelligibility and blindness were both non-significant (*p=*.51, *p*=.377). Follow-up post-hoc one-sample t-tests on the phase slope estimates for each group individually revealed a significant direction bias from CS to STG for SI (*t*(14)=-2.22, *p=.044*). No directional bias was found for EB (*t*(16)=.74, *p*=.47) As depicted in Fig 2 C-D, these results suggest that CS projects predominantly to STG in SI, whereas in EB this interaction is likely more balanced, and trending in the opposite direction.

## Discussion

We took advantage on the statistical and conceptual power offered by correlating magnetoencephalographic recordings with the envelope of naturalistic continuous speech to test if neuronal populations in the early blind’s occipital cortex synchronize to the temporal dynamics present in human speech. Using source localized MEG activity, we confirm and extend previous research (Scott et al., 2000; Davis and Johnsrude, 2003; Narain et al., 2003; Rodd et al., 2005, 2010; Okada et al., 2010; Peelle et al., 2010) by showing that temporal brain regions entrain to the natural rhythm of speech signal in both blind and sighted groups (Fig 1F). Strikingly, we show that following early visual deprivation occipito-parietal regions enhance their synchronization to the acoustic rhythm of speech (Fig 1G), independently of language content. That is, the region around the Precuneus/Cuneus area, seems to be involved in low-level processing of acoustic information alone. Previous studies have demonstrated that this area enhance its response to auditory information after early but not late visual deprivation (Collignon et al., 2013) and maintain such elevated response to sound even after sight-restoration early in life (Collignon et al., 2015). These studies did not find that this crossmodal response was related to a specific cognitive function but rather pointed to a more general response to sounds (Collignon et al., 2013, 2015). Interestingly, the current data suggest that this area is capable of parsing complex temporal patterns in sounds but is however insensitive to the intelligibility of speech, again suggesting a more general role in sound processing.

In order to isolate the brain regions modulated by speech comprehension, we identified those regions showing a greater activity for amplitude modulations that convey speech information compared to amplitude modulations that do not. In addition to the right STG observed in both groups (Fig 1H), the blind showed enhanced cerebro-acoustic coherence during intelligible speech at the lowest level of the occipital hierarchy, in the vicinity of calcarine sulcus (V1; Fig. 1I). This pattern of local encoding was accompanied by enhanced occipito-temporal connectivity during speech comprehension in EB compared to SI. While SI show the expected feed-forward projections from occipital to temporal regions (Lamme et al., 1998), EB show a more balanced connectivity profile, trending towards the reverse temporal to occipital direction. These findings support the idea of a reverse hierarchical model (Büchel, 2003) of the occipital cortex in EB where the regions typically coding for “low-level” visual features in the sighted (e.g. visual contrast or orientation) participate in higher-level function (e.g. speech intelligibility) in EB. Indeed, previous studies have found increased activity in the primary “visual” cortex of early blind people during Braille reading (Sadato et al., 1996; Burton et al.,2002, 2012), verbal memory and verb generation tasks (Amedi et al., 2003) and during auditory language-related processing (Bedny et al., 2011). In line with our results, activity in primary occipital regions in early blinds is stronger in a semantic versus a phonologic task (Burton et al., 2003) and vary as a function of syntactic and semantic complexity (Roder et al., 2002; Bedny et al., 2011; Lane et al., 2015). Moreover, repetitive transcranial magnetic stimulation (rTMS) over the occipital pole induces more semantic than phonologic errors in a verb-generation task in early blind people (Amedi 2003). However, our results however clearly suggest that the involvement of the occipital pole for language is not fully abstracted from sensory inputs as previously suggested (Bedny, 2017) since we show that occipital regions entrain to the envelope of speech and is enhanced by its intelligibility. Occipital participation in tracking the flow of speech in EB may constitute an adaptive strategy to boost perceptual sensitivity at informational peaks in language.

## Mechanistic origins of envelope tracking in occipital cortex

Although language is not the only cognitive process that selectively activate the occipital cortex of early blind people, it is arguably one of the most puzzling. The reason is that reorganization of other processes such auditory motion perception, and tactile object recognition appears to follow the topography of the functionally equivalent visual processes in the sighted brain (Ricciardi et al., 2007a; Amedi et al., 2010; Dormal et al., 2016). For example, the hMT+/V5 complex, typically involved in visual motion in the sighted, selectively processes auditory (Poirier et al., 2006; Dormal et al., 2016; Jiang et al., 2016) or tactile (Ricciardi et al., 2007b) motion in the blind. However, in the case of language, such recruitment is striking in light of the cognitive and evolutionary differences between vision and language (Bedny et al., 2011). This lead to the proposal that at birth, human cortical areas are cognitively pluripotent: capable of assuming a broad range of unrelated cognitive functions(Bedny, 2017). However, this argument resides on the presupposition that language has no computational relation with vision. But does this proposition reflect with what we know about the relation between the visual system and the classical language network? Rhythmic information in speech has a well-known language related surrogate in the visual domain, with the observation of lip movements (Lewkowicz and Hansen-Tift, 2012; Park et al., 2016). Indeed, both acoustic and visual speech signals exhibit rhythmic temporal patterns at prosodic and syllabic rates (Caplier et al., 2009; Schwartz and Savariaux, 2014; Giordano et al., 2017). The perception of lip kinematics that are naturally linked to amplitude fluctuations in speech serves as an important vehicle for the everyday use of language (Kuhl and Meltzoff, 1982; Weikum et al., 2007; Lewkowicz and Hansen-Tift, 2012) and help language understanding, especially in noisy conditions (Ross et al., 2007). Actually, reading lips in the absence of any sound activates both primary and association auditory regions overlapping with regions active during the actual perception of spoken words(Calvert et al., 1997). The synchronicity between auditory and visual speech entrain rhythmic activity in the observer’s primary auditory and visual regions and facilitate perception by aligning neural excitability with acoustic or visual speech features (Schroeder et al., 2008; Schroeder and Lakatos, 2009; Giraud and Poeppel, 2012; Mesgarani and Chang, 2012; Peelle and Davis, 2012; van Wassenhove, 2013; Zion Golumbic et al., 2013b; Park et al., 2016; Giordano et al., 2017). These results strongly suggest that both auditory and visual components of speech are processed together at the earliest level possible in neural circuitry based on the shared slow temporal modulations (around 2–7 Hz range) present across modalities (Caplier et al., 2009). Corroborating this idea, it has been demonstrated that neuronal populations in visual cortex follow the temporal dynamics of lip movements in sighted individuals, similar to the way temporal regions follow the acoustic and visual fluctuations of speech (Luo et al., 2010; Zion Golumbic et al., 2013a; Park et al., 2016). Like the temporal cortex, occipital cortex in the sighted also shows enhanced lip tracking when attention is directed to speech content. This result highlights the fact that a basic oscillatory architecture for tracking the dynamic aspects of (visual-) speech in occipital cortex exists even in sighted individuals.

Importantly, audiovisual integration of the temporal dynamics of speech has been suggested to play a key role when learning speech early in life: young infants detect, match and integrate the auditory and visual temporal coherence of speech (Kuhl and Meltzoff, 1982; Rosenblum et al., 1997; Lewkowicz, 2000, 2010; Brookes et al., 2001; Patterson and Werker,2003; Lewkowicz and Ghazanfar, 2006; Kushnerenko et al., 2008; Bristow et al., 2009; Pons et al., 2009; Vouloumanos et al., 2009; Lewkowicz et al., 2010; Nath et al., 2011). For instance, young infants between 4 and 6 months of age can detect their native language from lip movements only (Weikum et al., 2007). Around the same period, children detect synchrony between lip movements and speech sounds, and distribute more attention towards the mouth than towards the eyes (Lewkowicz and Hansen-Tift, 2012). Linking what they hear to the lip movements they observe may provide young infants with a stepping-stone towards language production (Lewkowicz and Hansen-Tift, 2012). Moreover, infants aged 10 weeks already exhibit a McGurk effect; highlighting again the early multisensory nature of speech perception (Rosenblum et al., 1997). Taken together, these results suggest that an audio-visual link between observing lip movements and hearing speech sounds is present at very early developmental stages, potentially from birth, which helps infants acquire their first language.

In the context of these evidences, the current results may support the biased connectivity hypothesis of cross-modal reorganization (Reich et al., 2011; Collignon et al., 2012; Hannagan et al., 2015; Striem-amit et al., 2015). Indeed, it has been argued that reorganization in blind occipital cortex might be constrained by functional pathways to other sensory- and cognitive systems that are also present in sighted individuals (Hannagan; Johnson). This hypothesis may explain the overlap in functional specialization between blind and sighted individuals (Collignon et al., 2011; Dormal et al., 2016; He et al., 2013; Jiang et al., 2016; Peelen et al., 2013; Pietrini et al., 2004; Poirier et al., 2006). In the context of our experiment, sensitivity to acoustic dynamics of intelligible speech in blind occipital cortex could arise from pre-existing occipito-temporal pathways connecting the auditory and visual system that are particularly important for the early developmental stages of language acquisition. In fact, the reorganization of these potentially pre-disposed pathway to process language content would explain how language-selective response may appear as early as in 3yo blind children (Bedny et al., 2015).

Previous studies have suggested that language processing in the occipital cortex arise through top-down projections from frontal region typically associated to the classical language network (Bedny et al., 2011; Deen et al., 2015) and that the representational content is symbolic and abstract, rather than sensory(Bedny, 2017). Our results contrasts with this view by showing that neuronal populations in (peri-) calcarine cortex align to the temporal dynamics of intelligible speech, and are functionally connected to areas sensitive to auditory information in temporal cortex. In sighted individuals, regions of the temporal lobe including STS are sensitive to acoustic features of speech, whereas higher-level regions such as anterior temporal cortex and left inferior frontal gyrus are relatively insensitive to these features and therefore do not entrain to the syllabic rate of speech (Davis and Johnsrude, 2003; Hickok and Poeppel, 2007; see confirmation in figure 1F-G). This suggests that occipital areas respond to speech at a much lower – sensory – level than previously thought in EB, potentially based on the reorganization of existing multisensory pathways connecting ‘auditory’ and ‘visual’ centres in the brain. Functional dependencies between sensory systems indeed exist between the earliest stages of sensory processing in humans (Ghazanfar and Schroeder, 2006; Kayser et al., 2008; Schroeder and Lakatos, 2009; Murray et al., 2016) and in nonhuman primates (Falchier et al., 2002; Lakatos et al.,2007; Schroeder and Lakatos, 2009)(Lakatos et al. 2005, 2007, suggesting an evolutionarily conserved mechanism of sensory processing. We therefore postulate that brain networks typically dedicated to the integration of audio-visual speech signal might be reorganized in the absence of visual inputs and lead to an extension (eg. lack of synaptic pruning) of speech tracking in the occipital cortex. Building on this connectivity bias, the occipital pole may extend its sensitivity to the intelligibility of speech, a computation this region is obviously not dedicated for in the first place.

Actually, a number of recent studies have suggested that visual deprivation reinforces the functional connections between the occipital cortex and auditory regions typically classified as the language network (Hasson et al., 2016; Schepers et al., 2012). Previous studies relying on dynamic causal modelling bring partial support to the idea that auditory information reaches the reorganized occipital cortex of the blind through direct temporo-occipital connection rather than using subcortical (Klinge et al., 2010) or top-down pathways (Collignon et al., 2013). In support of these studies, we observed that the overall magnitude of functional connectivity between occipital and temporal cortex is higher in the blind versus sighted individuals during natural speech comprehension. Moreover, directional connectivity analysis revealed that the interaction between the two cortices is also qualitatively different; sighted individuals show a strong feed forward drive towards temporal cortex, while blind individuals show a more balanced information flow, and a trend in the reverse direction. Our hypothesis that speech processing in the occipital cortex of EB arises from direct occipito-temporal connection is consistent with previous results showing that entrainment of theta activity in auditory cortex is insensitive to top-down influences from left inferior frontal regions (Kayser et al., 2015; Park et al., 2015).

## Speech comprehension in the right hemisphere

We observed that neuronal populations in right superior temporal cortex synchronize to the temporal dynamics of intelligible, rather than non-intelligible speech in both groups combined. Why does speech intelligibility modulate temporal regions of the right, and not the left, hemisphere? According to an influential model in speech comprehension, the asymmetric sampling in time model ( AST; Giraud and Poeppel, 2012; Hickok and Poeppel, 2007; Poeppel, 2003), there is a division of labour between left- and right auditory cortices (Poeppel, 2003; Boemio et al., 2005; Hickok and Poeppel, 2007), with left auditory cortex being more sensitive to high frequency information (+20Hz), whereas right temporal cortex responds mostly to low frequency information (~6Hz), such as syllable sampling and prosody (Belin et al., 1998; Poeppel, 2003; Boemio et al., 2005; Obleser et al., 2008; Giraud and Poeppel, 2012). Several studies have shown that the right hemisphere more specifically involves in the representation of connected speech (Bourguignon et al., 2013; Fonteneau et al., 2015; Horowitz-Kraus et al., 2015; Alexandrou et al., 2017), and previous studies directly demonstrated the prevalence of speech-to-brain entrainment in delta and theta bands in the right hemisphere more than the left during the listening of sentences or stories (Luo and Poeppel, 2007; Abrams et al., 2008; Gross et al., 2013; Giordano et al., 2017) The present study therefore replicates those results by showing enhanced phase coupling between the right hemisphere and the speech envelope at the syllabic rate (low-frequency phase of speech envelope), consistent with the AST-model. An interesting observation in the current study is that right hemispheric sensitivity to intelligible speech in temporal areas coincides with the enhanced right hemispheric sensitivity to intelligible speech in blind occipital cortex. An important open question for future research concerns the relative behavioral contribution of occipital and perisylvian cortex to speech understanding.

## Conclusion

We demonstrate that the right primary “visual” cortex synchronizes to the temporal dynamics of intelligible speech, at the rate of syllable transitions in language (~6-7Hz). These results demonstrate that the involvement of this neural population relates to the sensory signal of speech and therefore contrasts with the proposition that occipital involvement in speech processing is abstracted from its sensory input and purely reflect higher-level operations similar to those observed in prefrontal regions(Bedny, 2017). Blindness, due to the absence of organizing visual input, leaves the room open for sensory and functional colonization of occipital regions. This colonization will however not be stochastic, but will be constrained by modes of information processing natively anchored in specific brain regions and networks. Even if the exact processing mode has still to be unveiled by future research, we presume that language map onto occipital cortex building on pre-existing oscillatory architecture typically linking auditory and visual speech rhythm (Caplier et al., 2009). Our study therefore supports the view that the development of functional specialisation in the human cortex is the product of a dynamic interplay between genetic and environmental factors during development, rather than being predetermined at birth (Elman et al., 1996).

## Material and Methods

### Participants

Seventeen early blind (11 female, mean ± SD, 32.9 ± 10.19 years, range, 20-67 years) and sixteen sighted individuals (10 female, mean ±SD, 32.2 ± 9.92 years, range, 20-53 years) participated in the current study. There was no age difference between the blind and the sighted group (*t*(30) = .19, *p*= .85). All participants were either totally blind, or severely visually impaired from birth, however, two participants reported residual visual perception before the age of 3, one before the age of 4 and one participant lost sight completely at age 10. Causes of vision loss were damage to, or detached retina (10), damage to the optic nerve (3), infection ofthe eyes (1), microphtalmia (2), and hypoxia (1). While some participants reported residual light perception, none was able to use vision functionally. All participants were proficient braille readers, and native speakers of Italian. None of them suffered from a known neurological or peripheral auditory disease. The data of one sighted individual was not considered due to the discovery of a brain structural abnormality unknown to the experimenters at the time of the experiment. The project was approved by the local ethical committee at the University of Trento. In agreement with the Declaration of Helsinki, all participants provided written informed consent to participate in the study.

### Experimental design

Auditory stimuli were delivered into the magnetically shielded MEG room via stereo loudspeakers using a Panphonics Sound Shower 2 amplifier at a comfortable sound level, which was the same for all participants. Stimulus presentation was controlled via the Matlab Psychophysics Toolbox 3 [http://psychtoolbox.org] running on a Dell Alienware Aurora PC under Windows 7 (64 bit). Both sighted and blind participants were blindfolded during the experiment, and the room was dimly lit to allow for visual monitoring of the participant via a video feed from the MEG room. Instructions were provided throughout the experiment using previous recordings from one of the experimenters (FB).

The stimuli consisted of 14 short segments (~1min) from popular audiobooks (e.g., Pippi Longstocking and Candide) in Italian (nat). Furthermore, channel-vocoding in the Praat software was used to produce two additional control conditions. First, the original sound file was band pass filtered into 1 (1-channel) or 8 (8-channel) logarithmically spaced frequency bands. The envelope for each of these bands was computed, filled with Gaussian white noise, and the different signals were recombined into a single sound file. The resulting signal has an amplitude envelope close to the original, while the fine spectral detail is gradually distorted. Perceptually, the voice of the speaker is highly distorted in the 8-channel condition, however intelligibility is unperturbed. In contrast, in the 1-channel condition, speech is entirely unintelligible. In total, 42 sound files were presented in a pseudo-randomized fashion, distributed among seven blocks. Each block contained two sound files from each condition, and the same story was never used twice in the same block. Examples of the stimuli are provided in supplementary media content (S1).

To verify story comprehension, each speech segment was followed by a single-sentence statement about the story. Participants were instructed to listen to the story carefully, and judge whether the statement at the end was true or false, using a nonmagnetic button box. Responses were provided with the index and middle finger of the right hand.

### MEG data acquisition and preprocessing

MEG was recorded continuously from a 306 triple sensor (204 planar gradiometers; 102 magnetometers) whole-head system(Elekta Neuromag, Helsinki, Finland) using a sampling rate of 1kHz and online band-bass filter between 0.1 and 300Hz. The headshape of each individual participant was measured using a Polhemus FASTRAK 3D digitizer. Head position of the subject was recorded continuously using five localization coils (forehead, mastoids).

Data pre-processing was performed using the open-source Matlab toolbox Fieldtrip [www.fieldtriptoolbox.org], as well as custom code. First the continuous data was filtered (high-pass Butterworth filter at 1Hz, low-pass Butterworth filter at 170-Hz, and DFT filter at 50,100, and 150Hz to remove line-noise artefacts in the signal), and downsampled to 256Hz. Next, the data were epoched into segments of 1 second for subsequent analysis.

Rejection of trials containing artefacts and bad channels was performed using a semi-automatic procedure. First, outliers were rejected using a pre-screening based on the variance and range in each trial/channel. Then, algorithmically guided visual inspection of the raw data was performed to remove any remaining sources of noise.

### Extraction of the speech amplitude envelope

Amplitude envelopes of the stories were computed using the Chimera toolbox (Caplier et al., 2009) and custom code, following the procedure described by Gross and colleagues (Gross et al., 2013). First, the sound files were band-pass filtered between 100 and 1000Hz into 9 frequency-bands, using a 4^th^ order Butterworth filter. The filter was applied in forward and backward direction to avoid any phase shifts with respect to the original signal, and frequency-bands were spaced with equal width along the human basilar membrane. Subsequently, the analytic amplitude for each filtered segment was computed as the absolute of the Hilbert transform. Finally, the amplitude envelopes of all 9 bands were summed and scaled to a maximum value of 1. The resulting envelope was combined with the MEG data, and processed identically henceforth.

### Analysis of cerebro-acoustic coherence

To determine where, and at what temporal scale neuronal populations follow the temporal dynamics of speech, we computed spectral coherence between the speech envelope and the MEG signal. Coherence is a statistic that quantifies the phase relationship between two signals, and can be used to relate oscillatory activity in the brain with a peripheral measure such as a speech signal (Peelle et al., 2013). The first analysis was conducted in sensor space, across conditions and participants, to determine at what temporal scale coherence between the speech envelope and the MEG signal is strongest. To this end, a Hanning taper was applied to the 1s data segments, and Fourier transformation was used to compute the cross-spectral density between 1 and 30Hz, with a step size of 1Hz. To render the coherence values more normally distributed, a Fisher z-transform was applied by computing the inverse hyperbolic tangent (atanh).

Source-space analysis was centred on the coherence frequency-band of interest (FOI), identified in the sensor space analysis. Source reconstruction was performed using a frequency domain beamformer called Dynamic Imaging of Coherent Sources (DICS) (Gross et al., 2001; Liljeström et al., 2005). DICS was used to compute cerebro-acoustic coherence for all locations on a 3-dimensional grid (8x8x8mm). The forward model was based on a realistic single-shell headmodel(Nolte, 2003) for each individual. As structural scans could not be acquired for all participants, we approximated individual anatomy by warping a structural MNI template brain (MNI, Montreal, Quebec, Canada; www.bic.mni.mcgill.ca/brainweb) into individual headspace using the information from each participant’s headshape.

### Source connectivity analysis

Functional connectivity between occipital and temporal cortex was computed by extracting virtual sensor time-series at the locations of interest in CS and STG using time-domain beamforming. These virtual sensor time series were used to compute non-directional and directional connectivity metrics.

Virtual sensor time-series at the two locations of interest were extracted using a time-domain vector-beamforming technique called linear constrained minimum variance (LCMV) beamforming (Van Veen et al., 1997). First, average covariance matrices were computed for each participant to estimate common spatial filter coefficients. These filter coefficients were multiplied with the single-trial cleaned MEG data. To reduce the resulting three-dimensional time-series to one, singular value decomposition (SVD) was applied, resulting in a single time-series for each trial and participant.

Functional connectivity between the two regions was computed at peak locations using the phase-locking value (PLV) (Lachaux et al., 1999). First, single-trial virtual sensor time courses were converted to the frequency domain (0-50Hz) using Fourier transformation. The data was padded to 2s and a Hanning taper was applied to reduce spectral leakage. Coherence was used as a proxy for functional connectivity. To disentangle phase consistency between regions from joint fluctuations in power, the spectra were normalized with respect to the amplitude, resulting in an estimate of the phase-locking (Lachaux et al., 1999) between regions. Connectivity estimates were normalized using a Fischer z-transform (atanh), as in the analysis of cerebro-acoustic coherence.

PLV is a symmetric proxy for connectivity and does not allow for inferences regarding the direction of information flow between two regions of interest. To test whether differences in functional connectivity between EB and SI are accompanied by changes in the directionality of the flow of information between the regions of interest, we computed the phase slope index (PSI) (Nolte et al., 2008). The phase slope index deduces net directionality from the time delay between twotime series (x1 and x2). Timeseries x1 is said to precede, and hence drive timeseries x2, in a given frequency band, if the phase difference between ×1 and ×2 increases with higher frequencies. Consequently, a negative phase slope reflects a net information flux in the reverse direction, that is, from ×2 to ×1. Here, we computed the phase slope index using a bandwidth of ±5Hz around the frequencies of interest. Following the recommendation by Nolte and colleagues (Nolte et al., 2008), PSI estimates were normalized with the standard error, which was computed using the jackknife method.

### Statistical analysis

Statistical testing of the behavioural comprehension scores as well as the connectivity estimates was performed using linear mixed-effects models in R. Differences between conditions and groups in source space were evaluated using Statistical Parametric Mapping (SPM12), and the Threshold-Free Cluster Enhancement (TFCE) toolboxes in Matlab. TFCE (Smith and Nichols, 2009) computes new values for each voxel in a statistical map as a function of the original voxel value and the values of the surrounding voxels. By enhancing the statistical values in voxels with a high T-value that are also part of a local cluster, TFCE optimally combines the benefits of voxel-wise and cluster-based methods. TFCE was applied to the whole-brain contrasts. Final correction for multiple comparisons was applied using a maximum statistic based on 1000 permutations of group membership (independent testing) or conditions (dependent testing). Here, we applied a variance smoothing of 15mm FWHM to reduce the effects of high spatial frequency noise in the statistical maps. The smoothing kernel used in the current study is higher than in comparable fMRI studies due to the inherent smoothness of the source-reconstructed MEG data.

## Acknowledgments

The project was funded by the ERC grant MADVIS – Mapping the Deprived Visual System: Cracking function for prediction (Project: 337573, ERC-20130StG) awarded to Olivier Collignon. We would also like extend our gratitude to Valeria Occelli, and the Master students who have assisted with the data acquisition: Marco Barilari, Giorgia Bertonati, and Elisa Crestale for their kind assistance with the blind participants. In addition, we would like to thank Gianpiero Monittola for ongoing support with the hardware. Finally, we would like to thank the organizations for the blind in Trento, Mantova, Genova, Savona, Cuneo, Torino, Trieste and Milano.

## Author Contributions

MJVA, OC, and FB designed the experiment. MJVA, FB, RB, and SM screened and tested the participants. MJVA analysed the data. MJVA and OC wrote the manuscript. FB, RB and SM did comments on the draft of the manuscript.

## Supplementary materials

S1. Movie of the natural and two conditions

